# Small extracellular vesicles from young adipose-derived stem cells ameliorate age-related changes in the heart of old mice

**DOI:** 10.1101/2024.06.06.597840

**Authors:** Jorge Sanz-Ros, Javier Huete-Acevedo, Cristina Mas-Bargues, Nekane Romero-García, Mar Dromant, Michel van Weeghel, Georges E. Janssens, Consuelo Borrás

## Abstract

**Background:** Aging is a process characterized by a progressive decline in physiological functions. This decline contributes to an increased risk of various age-related diseases, including conditions such as heart failure or aortic stenosis. Recent advances have highlighted the potential of stem cell therapies in mitigating the adverse effects of aging. The mechanisms underlying the therapeutic effects of these cells appear to be largely paracrine, including the exchange of different molecules via extracellular vesicles (EVs). This work aims to investigate the potential of sEVs derived from young adipose-derived stem cells (ADSC-sEVs) in restoring structural, molecular, and functional changes associated with aging in the heart.

**Methods:** Aged C57BL/6J mice were treated intravenously with ADSC-sEVs from young mice or PBS as controls, young mice were also included to identify specific changes associated with aging. The impact of sEV treatment on cardiac function was assessed using transthoracic echocardiography and physical endurance tests. Histological and molecular analyses were conducted on heart tissue to evaluate structural changes and markers of senescence, inflammation, and oxidative stress. Additionally, a comprehensive metabolomic analysis was performed on heart tissues to identify changes in metabolic profiles associated with aging and treatment status.

**Results:** Administration of ADSC-sEVs significantly improves several aging-associated cardiac parameters, including reductions in oxidative stress, inflammation, and cellular senescence. We also report on age-related changes reversal in myocardial structure and function, highlighted by decreased fibrosis and improved vascularization. Notably, echocardiographic assessments reveal that sEV treatments ameliorate diastolic dysfunction and left ventricle structural alterations typically associated with aging. Furthermore, the treatment shifts the heart metabolome towards a more youthful profile.

**Conclusions:** These results denote the potential of ADSC-sEVs as a novel, non-invasive therapeutic strategy to mitigate cardiac aging-associated functional decline.

## Introduction

Aging is accompanied by a decline in tissue regenerative capacity, leading to an increased susceptibility to various age-related diseases^1,2^. While stem cell therapies have shown promise in regenerative medicine, recent studies suggest that their therapeutic effects are predominantly mediated by paracrine mechanisms rather than direct tissue integration^3–5^. Extracellular vesicles (EVs), including small extracellular vesicles (sEVs), have emerged as crucial mediators of intercellular communication, carrying a diverse cargo of proteins, nucleic acids, and lipids^6,7^.

Mesenchymal stem cells (MSCs) have been extensively investigated for their regenerative potential due to easy isolation, culture, and low immunogenicity^8^. However, the use of MSCs is hindered by their limited survival and integration into host tissues. In contrast, the secretome of MSCs, particularly EVs, offers advantages such as stability, ease of dosage, and lower immunogenicity^9^, with some clinical trials for several conditions ongoing^10,11^. EVs are also being explored as regulators of aging-related processes, participating in cellular senescence, oxidative stress, telomere dysfunction, inflammation, or metabolic dysregulation^12,13^.

The human average lifespan is on the rise, resulting in a significant increase in the population aged 65 and older, a trend expected to persist over the next two decades^14^. Within this demographic, cardiovascular disease remains the primary cause of mortality, accompanied by escalating treatment costs^15^. Aging, an inherent aspect of life, stands as the predominant risk factor for cardiovascular diseases^16^. The relationship between cardiovascular disease and the molecular and cellular aspects of aging is therefore clear. Heart tissue suffers several changes associated with aging, such as fibrosis, valvular degeneration, and calcification^17^, cardiomyocyte hypertrophy^18^, degeneration of the conduction system^19^, or cellular senescence-associated changes^20^. These changes contribute to the development of several diseases associated with old age, whose prevalence and incidence are rising, such as heart failure with preserved ejection fraction (HFpEF) or aortic stenosis^21^.

The present study focuses on the therapeutic potential of ADSC-sEVs in mitigating age-related changes in the heart. Here we show that ADSC-sEVs exert anti-aging effects in the heart of old mice through the modulation of cellular senescence, oxidative stress, inflammation, and metabolism.

## Material and methods

### Mouse model

This investigation adhered rigorously to all relevant federal and institutional guidelines. The University of Valencia Animal Ethics Committee, following European Union (EU) regulations on animal research, approved the protocol under identification numbers A1490612630929 and A1508582840889. The mice utilized in the study were of a C57BL/6J background, with aged mice treated at 20 to 24 months. Both genders were included, and, when feasible, littermates of the same sex were used. The sample size was determined using G*Power, and random assignment to experimental groups was conducted. Each mice received two doses separated by 7 days, comprising either 20 μg of sEV protein or PBS as a control, the doses were administered intravenously to mice via the tail vein in a total volume of 100 μl diluted in PBS. The protocol was replicated across several batches of old mice treated with sEVs versus PBS as a control. Mouse weight was annotated during the experimental procedure as a surrogate marker of toxicity. Upon euthanasia 30 days post-treatment with sEVs/PBS, the heart was collected, weighed, and stored at -80°C for further processing. Please see the Major Resources Table in the Supplemental Materials.

### Stem cell isolation and culture

Stem cell culture involved obtaining mesenchymal stem cells (MSCs) from inguinal fat pads of mice aged 3 to 6 months (ADSCs) according to the previously described protocol^22^. Cells in passages 2 or 3 were used, cultured in high-glucose Dulbecco’s modified Eagle’s medium (DMEM) with 10% FBS and 1% penicillin/streptomycin. Culture conditions included 37°C, 5% CO2, and 3% O2. ADSCs’ characterization was conducted as referenced in our previous work^23^.

### Isolation of ADSC-derived sEVs

Isolation of ADSC-derived small extracellular vesicles (sEVs) involved culturing ADSCs in high-glucose DMEM with 2% exosome-depleted FBS and 1% penicillin/streptomycin (Gibco, A2720803) for 48 hours. Then, the conditioned medium was collected, and sEVs were isolated by differential ultracentrifugation. The culture medium underwent centrifugation at 2000g for 10 minutes and subsequently at 20,000g for 30 minutes to eliminate whole cells, cell debris, and larger extracellular vesicles (EVs). The resulting supernatant underwent ultracentrifugation at 100,000g for 70 minutes. The pelleted vesicles were then re-suspended in PBS, subjected to another round of ultracentrifugation at 100,000g for 70 minutes for thorough washing, and re-suspended in PBS again. sEVs isolated from the conditioned medium were stored at 4°C and employed for treatment within a 24-hour timeframe. sEV dosage was determined by protein quantification using the Lowry method.

### Transmission electron microscopy and immunogold labelling of sEVs

The isolated sEVs were fixed in a solution of 2% paraformaldehyde (PFA) in 0.1 M PBS for 30 minutes. A glow discharge technique (30 seconds, 7.2 V) using a Bal-Tec MED 020 Coating System was applied to carbon-coated copper grids. Subsequently, these treated grids were positioned over the sample drops for 15 minutes. The grids with attached sEVs were then immersed in a 0.1 M PBS solution for washing, followed by an additional fixation step in 1% glutaraldehyde for 5 minutes. After thorough washing in distilled water, the grids were contrasted with 1% uranyl acetate and embedded in methylcellulose. Excess fluid was carefully removed, and the samples were allowed to dry before examination using a transmission electron microscope FEI Tecnai G2 Spirit (Thermo Fisher Scientific, OR, USA). Image acquisition was performed using a digital camera Morada (EMSIS GmbH, Münster, Germany). For immunogold labelling, 8 μl of isolated sEVs underwent fixation in 2% PFA in 0.1 M PBS for 30 minutes. Carbon-coated nickel grids were then placed over these sEV drops for 15 minutes. After washing in 0.1 M PBS, the grids were blocked in a solution of 0.1 M glycine and 0.3% BSA for 10 minutes. The grids were subsequently incubated with the primary antibody anti-CD63 (MBL, D263-3) at a dilution of 1:100 for 1 hour. Following another blocking step for 10 minutes, the grids were exposed to Gold 6nm–conjugated goat anti-rat secondary antibody (Abcam, ab105300) at a dilution of 1:1000 for 1 hour. Lastly, after thorough washing, a standard negative staining procedure was employed, and the samples were observed under a transmission electron microscope, as previously described.

### PKH26 staining

For *in vivo* tracking of sEVs, we obtained sEVs from the conditioned medium of young ADSCs. Subsequently, these sEVs were labeled with 5 μM PKH26 (Sigma-Aldrich, MINI26-1KT) following the first ultracentrifugation. Following the labeling, the sEVs were resuspended in PBS and subjected to an additional round of washing through ultracentrifugation. A total of 20 µg of PKH26-labeled sEVs in 100 µL of PBS were injected into aged mice (20 months), control mice were injected 100 µL of a PBS solution containing 5 μM PKH26. The mice were euthanized 24 hours later, and heart tissue was harvested and fixed in 4% PFA for 24 hours. 50-micron slices were prepared, stained with DAPI (Invitrogen) at a 1:1000 dilution for 30 minutes at room temperature, and mounted on coverslips using an aqueous mounting medium sealed with nail polish. Imaging was conducted using an Olympus FV1000 confocal laser scanning biological microscope. Images were processed in ImageJ maintaining equal ratios.

### Echocardiography assessment in mice

Anesthesia was induced in a chamber with 2-3% isoflurane delivered in a mixture of oxygen (1-2 L/min). Once anesthetized, mice were transferred to a nose cone for continuous isoflurane inhalation at a maintenance concentration of 1-2%. Anesthesia depth was monitored throughout the procedure by assessing the respiratory rate and pedal withdrawal reflex. Echocardiography was performed using an imaging system (GE Versana Active Veterinary Ultrasound Scanner) equipped with a 12-MHz transducer for small animals. Mice were positioned in a supine position with external heating to maintain body temperature. The chest area was shaved, and ultrasound gel was applied for optimal acoustic coupling. Two-dimensional (2D) M-mode and pulsed Doppler images were acquired in various views, including parasternal long-axis, short-axis, and apical views. For data analysis, images were analyzed in the same ultrasound scanner used for acquisition. Standard echocardiographic parameters shown in the results section were measured, for fractional shortening (FS) calculation, M-mode images were used to measure left ventricle end-diastolic diameter (LVEDD), and end-systolic diameter (LVESD), FS=LVEDD – LVESD / LVEDD x 100.

### Treadmill physical endurance test

The animals underwent a progressive intensity treadmill test (Treadmill Control LE 8710 Panlab, Harvard Apparatus) to assess their endurance, measured by maximum time running. Following a warm-up period, the treadmill belt velocity was incrementally elevated until the animals reached exhaustion, indicating their inability to continue running. The test commenced with an initial 4-minute session at 10 cm/s, followed by successive increments of 4 cm/s every 2 minutes. Exhaustion was defined as the point at which a mouse remained on the shock grid for 5 seconds instead of actively running.

### Histology

Heart tissue was freshly frozen in liquid nitrogen and 10-micron slices were obtained using a cryostat for histological staining and immunofluorescence. Hematoxylin and eosin staining (Sigma-Aldrich, MHS32, and E4009, respectively) or Sirius red staining (Sigma-Aldrich, 365548) was performed on the sections, followed by mounting and sealing for morphometric analysis. Images were captured using an optical microscope (Leica), and three images from different areas of each slice were obtained. Morphometric analysis of heart sections was conducted using ImageJ.

### Immunofluorescence imaging

Ten-micrometer tissue slices were mounted on slides and fixed with ice-cold acetone for 20 minutes. For permeabilization, slices were incubated in 1% Triton X-100 in PBS for 10 minutes. Sections were blocked with 10% normal goat serum (Invitrogen) in PBS with 0.05% Tween 20 (PBS-t) and incubated with primary antibodies overnight at 4°C in the same buffer. After primary antibody incubation, sections underwent three 10-minute PBS-t washes and were then incubated with secondary antibodies for 2 hours at room temperature. Subsequent washes were performed, and tissues were counterstained with DAPI (Invitrogen; 1:1000 dilution) for 30 minutes at room temperature. Coverslips were mounted with an aqueous mounting medium and sealed with nail polish. Images were acquired using an Olympus FV1000 confocal laser scanning biological microscope. Image processing was carried out using CellProfiler with custom pipelines for automatic cell counting and analysis of CD3, LMNB1, and γH2AX positive cells. CD31 area was calculated using ImageJ, with uniform adjustments to levels. Three images from different areas of each slice were obtained.

The following antibodies and concentrations were used:

CD31 staining: Alexa Fluor 647 anti-mouse CD31 antibody (Biolegend, 102515, 1:50 dilution). LMNB1 staining: anti-LMNB1 (Proteintech, 12987-1-AP; 1:50 dilution) and Alexa Fluor 488 anti-rabbit (Abcam, ab150077; 1:2000 dilution); a minimum of 150 nuclei were analyzed per sample.

γH2AX staining: anti-γH2AX (Cell Signaling Technology, 9718S; 1:1000 dilution) and Alexa Fluor 488 anti-rabbit (Abcam, ab150077; 1:2000 dilution); a minimum of 150 were analyzed per sample.

CD3 staining: anti-CD3 (Proteintech, 17617-1-AP; 1:1000 dilution) and Alexa Fluor 647 anti-rabbit (Abcam, ab150079, 1:2000 dilution).

### Quantification of lipid peroxidation by HPLC

Heart tissue was lysed using a KPi-EDTA buffer [50 mM KPi and 1 mM EDTA (pH 7.4)], and the levels of lipid peroxidation were assessed by quantifying malondialdehyde (MDA) using high-performance liquid chromatography (HPLC). MDA was determined as an MDA– thiobarbituric acid (TBA) adduct following a previously established method. This approach relies on the hydrolysis of lipoperoxides and the subsequent formation of a TBA-MDA2 adduct, which was detected using reverse-phase HPLC and quantified at 532 nm. The chromatographic technique was carried out under isocratic conditions, with the mobile phase comprising a mixture of monopotassium phosphate at 50 mM (pH 6.8) and acetonitrile (70:30). The MDA levels in each sample were normalized to the protein concentration determined by the Lowry method.

### Protein oxidation quantification with immunoblotting

Total protein carbonylation was detected by immunoblotting using the OxyBlot Protein Oxidation Detection kit (Merck) following the manufacturer’s instructions. In brief, tissues were lysed in tris/SDS/glycerol buffer, and protein concentration was determined using the Lowry method. Subsequently, 20 μg of proteins were separated on SDS polyacrylamide gels and transferred onto nitrocellulose membranes. The membranes were blocked with 3% BSA in PBS-t for 60 minutes at room temperature and then incubated overnight at 4°C with the primary antibody from the kit. After three washes (10 minutes each) with PBS-t, the membranes were incubated with the secondary antibody for 120 minutes at room temperature. Following three additional washes with PBS-t, the membranes were developed with Luminol (Sigma-Aldrich) using the ImageQuant LAS4000 system. Image analysis was performed in ImageJ, and Ponceau staining of the membranes served as the loading control.

### Interleukin quantification

We utilized two commercially available enzyme-linked immunosorbent assay (ELISA) kits for the quantitative analysis of IL-6 (Abcam, ab100713) and IL-8 (Abcam, ab234567), adhering to the manufacturer’s provided instructions. In brief, tissues underwent lysis using specific buffers included in each kit, protein concentration was determined using the Lowry method, and samples were appropriately diluted at ratios of 1:5. Absorbance at 450 nm was quantified using the Molecular Devices SPECTRAmax Plus 384. All samples were subjected to duplicate assays.

### Metabolomics

Metabolomics was performed as previously described, with minor adjustments^24,25^. A 75 µL mixture of the following internal standards in water was added to approximately 3 mg of freeze-dried heart tissue: adenosine-^15^N_5_-monophosphate (100 µM), adenosine-^15^N_5_-triphosphate (1 mM), D_4_-alanine (100 µM), D_7_-arginine (100 µM), D_3_-aspartic acid (100 µM), D_3_-carnitine (100 µM), D_4_-citric acid (100 µM), ^13^C_1_-citrulline (100 µM), ^13^C_6_-fructose-1,6-diphosphate (100 µM), guanosine-15N_5_-monophosphate (100 µM), guanosine-^15^N_5_-triphosphate (1 mM), ^13^C_6_-glucose (1 mM), ^13^C_6_-glucose-6-phosphate (100 µM), D_3_-glutamic acid (100 µM), D_5_-glutamine (100 µM), ^13^C_6_-isoleucine (100 µM), D_3_-leucine (100 µM), D_4_-lysine (100 µM), D_3_-methionine (100 µM), D_6_-ornithine (100 µM), D_5_-phenylalanine (100 µM), D_7_-proline (100 µM), ^13^C_3_-pyruvate (100 µM), D_3_-serine (100 µM), D_5_-tryptophan (100 µM), D_4_-tyrosine (100 µM), D_8_-valine (100 µM). Subsequently, 425 µL water, 500 µL methanol, and 1 mL chloroform were added to the same 2 mL tube before thorough mixing and centrifugation for 10 min at 14.000 rpm. The top layer, containing the polar phase, was transferred to a new 1.5 mL tube and dried using a vacuum concentrator at 60°C. Dried samples were reconstituted in 100 µL methanol/water (6/4; v/v). Metabolites were analyzed using a Waters Acquity ultra-high-performance liquid chromatography system coupled to a Bruker Impact II™ Ultra-High Resolution Qq-Time-Of-Flight mass spectrometer. Samples were kept at 12°C during analysis and 5 µL of each sample was injected. Chromatographic separation was achieved using a Merck Millipore SeQuant ZIC-cHILIC column (PEEK 100 x 2.1 mm, 3 µm particle size). Column temperature was held at 30°C. The mobile phase consisted of (A) 1:9 acetonitrile:water and (B) 9:1 acetonitrile:water, both containing 5 mM ammonium acetate. Using a flow rate of 0.25 mL/min, the LC gradient consisted of: Dwell at 100% Solvent B, 0–2 min; Ramp to 54% Solvent B at 13.5 min; Ramp to 0% Solvent B at 13.51 min; Dwell at 0% Solvent B, 13.51–19 min; Ramp to 100% B at 19.01 min; Dwell at 100% Solvent B, 19.01–19.5 min. Equilibrate the column using a 0.4 mL/min flow at 100% B from 19.5-21 min. MS data were acquired using negative and positive ionization in full scan mode over the range of m/z 50-1200. Data were analysed using Bruker TASQ software version 2021b (2021.1.2.452). All reported metabolite intensities were normalized to dry tissue weight, as well as to internal standards with comparable retention times and response in the MS. Metabolite identification was based on a combination of accurate mass, (relative) retention times, and fragmentation spectra, compared with the analysis of a library of standards. Statistical analysis and visualization of the acquired data were done in an R environment using the ggplot2, ropls, and mixOmics packages^26,27^.

### Statistical analysis

Ratios depicting the comparison of physical test values after treatment against baseline were calculated and presented as percentages over baseline, with the baseline defined as 0%. Outliers were assessed in all groups using the ROUT method (Q = 2%), with no exclusion of any data point. The normality of each group was assessed using the Shapiro-Wilk test. For pairwise comparisons, either the Unpaired Student’s t-test or the Mann-Whitney test was employed, depending on the data distribution. In the case of multiple comparisons, ANOVA was applied, with Tukey’s multiple comparisons serving as a post hoc test. Alternatively, for nonparametric data, the Kruskal-Wallis test was utilized, with Dunn’s multiple comparisons as the post hoc test. Each data point presented in the manuscript represents a biological replicate.. GraphPad Prism 9.0 software was utilized for both analysis and graphical design if not otherwise stated.

## Results

### Intravenously delivered sEVs from young ADSCs effectively reach the heart of old mice

Firstly, we conducted a characterization of the vesicles isolated from the cell culture supernatants of young ADSCs (Fig1), according to the minimal information for studies of extracellular vesicles (MISEV) recommendations for the characterization and functional studies of sEVs^7^. We utilized transmission electron microscopy (TEM) to evaluate the size and morphology of the isolated vesicles, revealing round-shaped vesicles ranging from 50 to 200 nm in diameter (Fig1). Additionally, the presence of CD63, a classical marker of sEVs, in the vesicle membrane was corroborated through immunogold labeling (Fig1). To demonstrate the delivery and uptake of sEVs in the heart tissue, we labeled the sEVs with a lipophilic dye (PKH26) and administered them into the bloodstream of old mice through tail vein injection. Subsequent histological study with fluorescence microscopy of the heart tissue revealed the uptake of the dye in this organ (Fig1).

**Figure 1:**
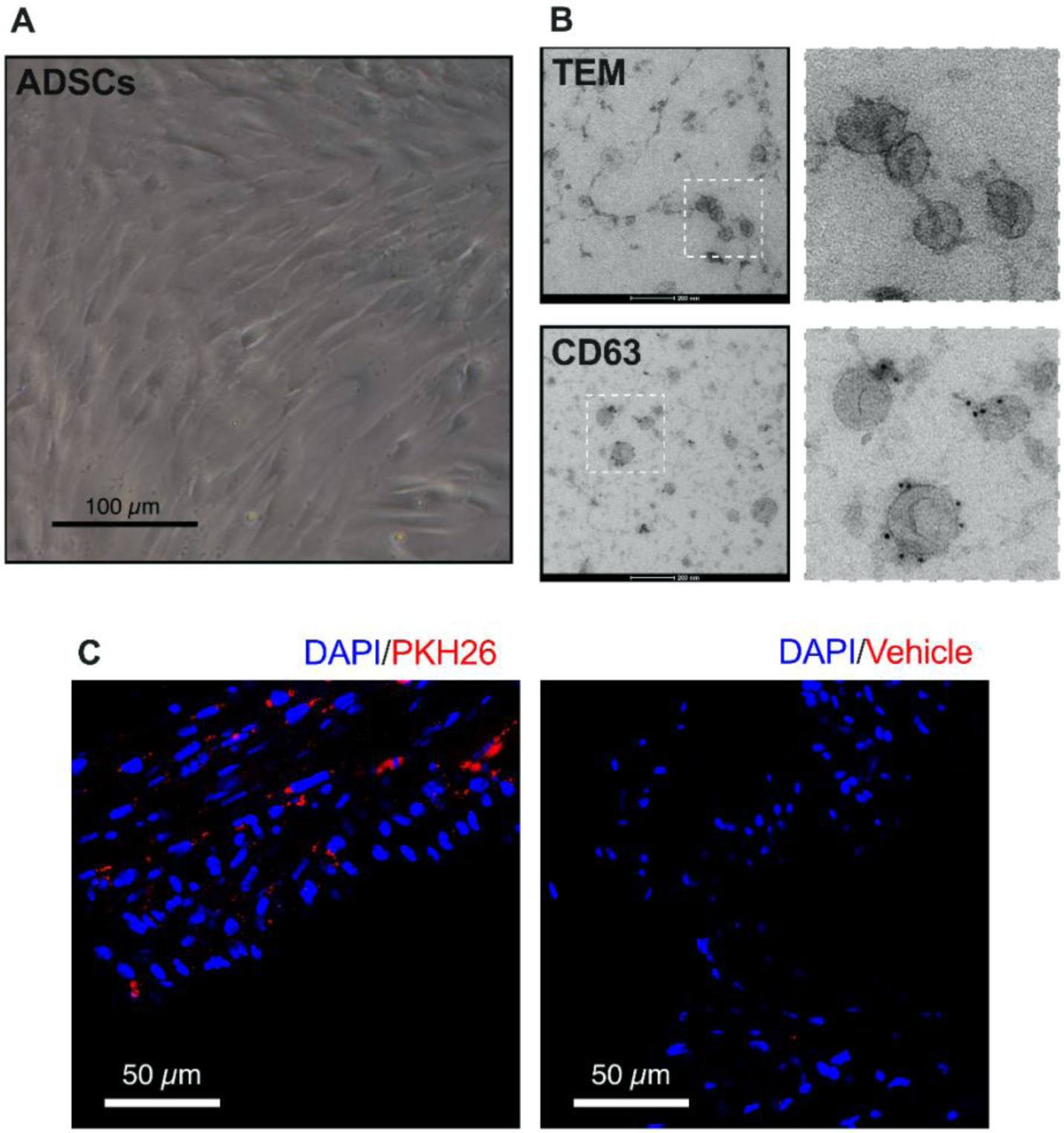
ADSC-sEVs reach the heart of old mice. (A) Representative image of passage 2 ADSCs in culture. (B) Representative images of transmission electron microscopy of ADSC-sEVs and immuno-gold labeling of the CD63 protein in sEVs samples. (C) Representative image of heart tissue from old mice treated with either PKH26 labeled ADSC-sEVs or vehicle as controls.

### ADSC-sEVs improve functional parameters associated with aging in the heart of old mice

We then assessed the impact of ADSC-sEVs on functional parameters associated with cardiac aging in old mice. To this end, we performed transthoracic echocardiography with a transducer designed for small animals in young (3-6 months) and old C57BL/6J mice (20-24 months), to obtain which parameters are altered with age in this mouse strain. To better characterize the effect of sEVs on echocardiographic parameters in old mice, we performed echocardiography before treatment (Day 0) and 30 days after treatment (Day 30) with PBS or sEVs (Fig2A).

**Figure 2:**
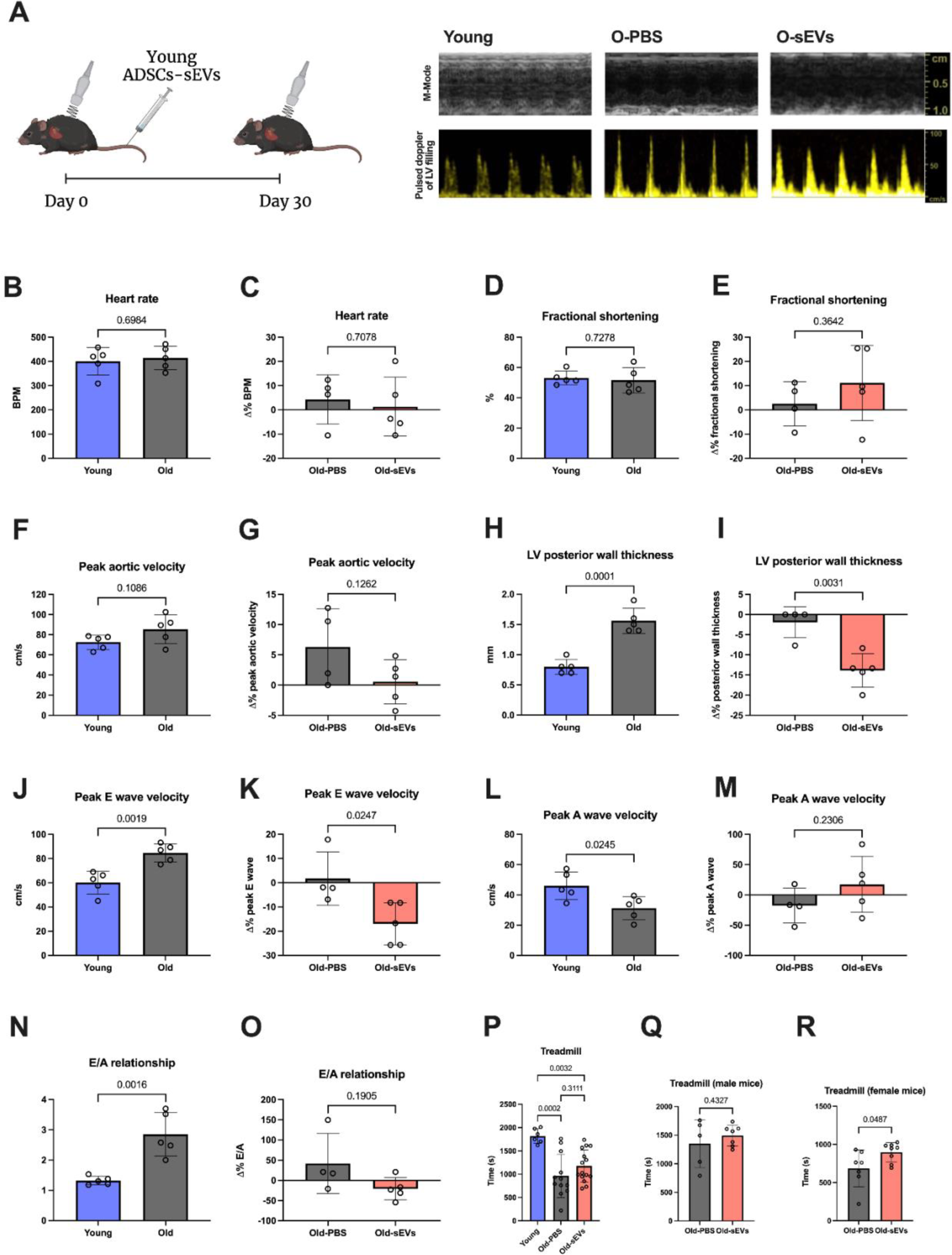
Effect of ADSC-sEVs on echocardiographic and functional parameters of old mice. (A) Schematic representation of the experimental design and representative images of M-mode and pulsed Doppler from transthoracic echocardiography. (B-C) Quantification of the heart rate. (D-E) Quantification of the fractional shortening as measured from LVEDD and LVESD measurements. (F-G) Quantification of peak aortic velocity from pulsed Doppler. (H-I) Quantification of LV posterior wall thickness. (J-K) Quantification of peak E wave velocity from pulsed Doppler of LV filling. (L-M) Quantification of peak A wave velocity from pulsed Doppler of LV filling. (N-O) E/A wave ratio measurement. (P-R) Quantification of maximum time running in the treadmill test, computed values and separated by sex. For young vs old comparisons, absolute values are shown. For O-PBS vs O-sEVs echocardiography comparisons, data are shown as the change in percentage from the baseline before treatment with sEVs/PBS. All data are shown as means ± SD. B-O: n=5 per group. P: n=6 for young, 12 for Old-PBS, 15 for Old-sEVs. Q: n=5 for Old-PBS, 7 for Old-sEVs. R: n=7 for Old-PBS, 8 for Old-sEVs.

Heart rate exhibited no significant changes from young to old mice, and this pattern persisted in old mice following sEVs treatment (Fig2B-C). Fractional shortening remained unaltered across age groups and treatment status, indicating little changes in systolic function during mouse aging, in line with previous studies^28^ (Fig2D-E). Peak aortic velocity displayed a tendency to increase in old mice, probably suggesting a propensity to develop aortic valve calcification in these mice, as previously noted^28,29^. sEVs treatment didn’t demonstrate a significant reduction in this trend (Fig2F-G).

The most prominent functional changes associated with aging in the heart, both in mice and humans, are the increase in left ventricle (LV) mass and reduced diastolic function^21,30^. Indeed, we found that LV posterior wall thickness exhibited an age-related increase in old mice compared to the young group; however, treatment with sEVs reversed this trend, resulting in a reduction in wall thickness (Fig2H-I). Regarding the pulsed Doppler of the LV filling, peak E wave velocity showed an increase with age in mice, conversely, peak A wave velocity decreased with age, leading to an increased E/A relationship (>2) in the old mice. This is indicative of an altered diastolic function with a restrictive pattern, where LV filling depends mainly on the protodiastolic wave. Treatment with sEVs in old mice partially reversed this pattern, with a significant reduction of the peak E wave velocity and a non-significant trend showing reduced E/A ratios (Fig2J-O).

As a measure of cardiac function, we performed a treadmill test to measure physical endurance in mice. Young mice showed higher levels of performance in the treadmill test compared to old mice (Fig2P). When adjusting for sex, female mice treated with sEVs showed an increased endurance in this test when compared to old female control mice (Fig2Q), which was not the case for male mice (Fig2R), suggesting a sex-dependent effect of the treatment, something common to diverse anti-aging interventions^31,32^.

Collectively, these findings show the potential of ADSC-sEVs in ameliorating age-associated alterations in cardiac functional parameters.

### ADSCs-sEVs alleviate age-related histological alterations of the mouse heart

To further correlate the effect of sEVs on the functional parameters, we explored the structural alterations in the heart tissue at the histological level. Some of these changes include the increase in LV mass, myocardial fibrosis, and alteration of the vascularization, all of which contribute to the functional decline, specially to the diastolic dysfunction observed during aging^29,33,34^.

In the first place, we measured the weight of the heart normalized by the total body weight (TBW) of the mice as an indicator of total heart mass. Old mice treated with sEVs showed a lower heart weight ratio when compared to controls (Fig3A), indicating a lower heart mass, the comparison between young and old mice was non-significant, probably because TBW is much lower in young mice. Histological analysis with Sirius red staining demonstrated an increase of fibrotic tissue in the hearts of old mice compared to young ones. Treatment with ADSCs-sEVs induced a significant decrease in fibrotic tissue (Fig3B and D), suggesting a protective effect against aging-associated cardiac remodeling and fibrosis. Furthermore, assessment of vascularization through CD31 immunostaining unveiled a significant reduction of the CD31+ area in the aged heart, while we observed an increase in the CD31+ area in heart slices from mice treated with ADSC-sEVs (Fig3C and E), indicating a pro-angiogenic effect of sEVs. This EV-dependent enhancement in vascularization has also been observed previously in several models of tissue damage^35,36^.

**Figure 3:**
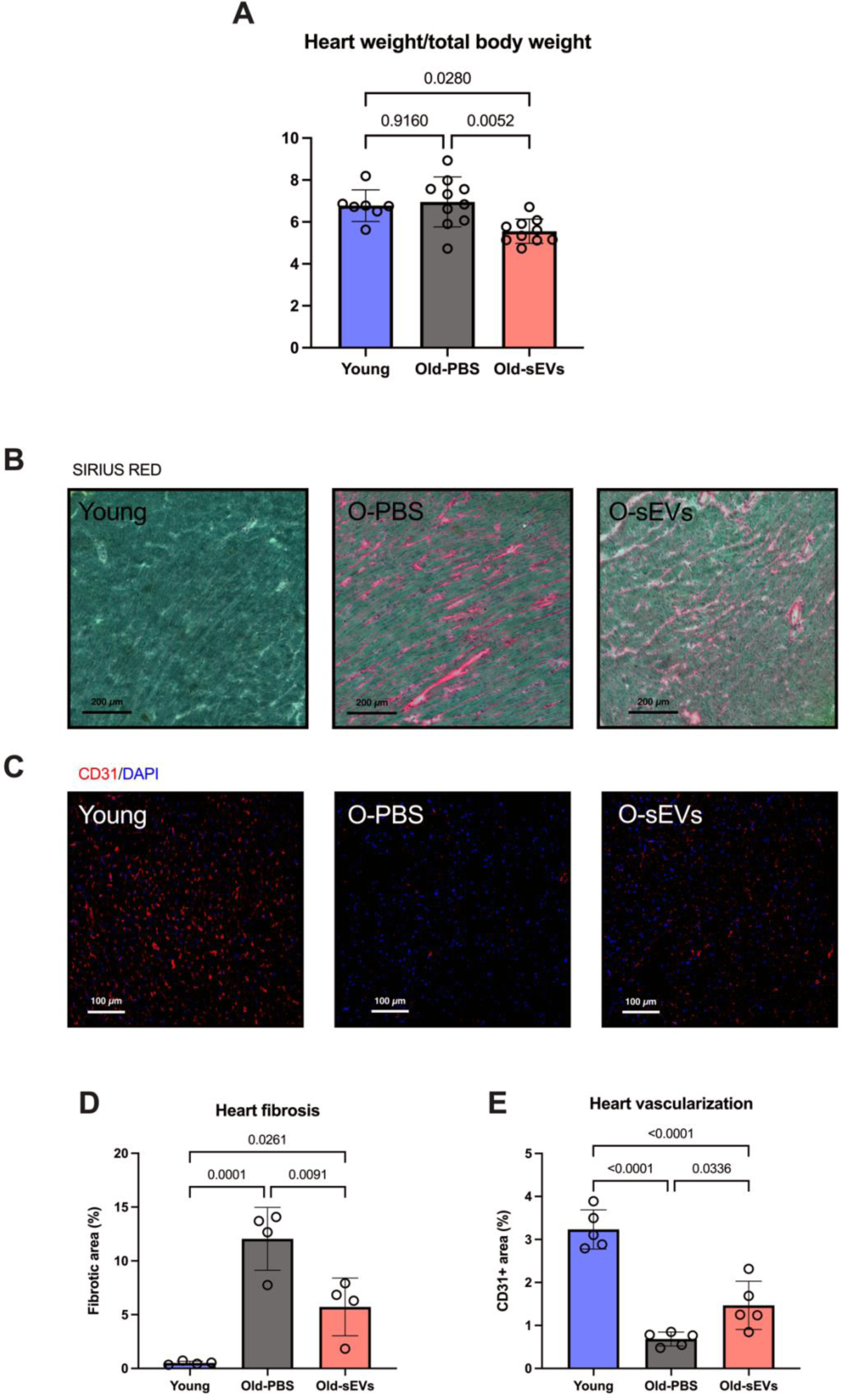
ADSC-sEVs mitigate structural changes in the mouse heart associated with aging. (A) Quantification of the ratio of heart weight divided by TBW. (B and D) Representative images and quantification of Sirius red staining in heart tissue under light microscopy. (C and E) Representative images and quantification of CD31 immunofluorescence staining of heart tissue under confocal microscopy. All data are shown as means ± SD. A: n=7 for young, 10 for Old-PBS and Old-sEVs. D: n=4 per group. E: n=5 per group.

These results show the role of ADSC-sEVs in counteracting age-related cardiac structural changes in aged mice.

### Cellular and molecular markers of aging are partially reversed by young ADSCs-sEVs in the heart of old mice

Some of the molecular and cellular markers of aging are common to the different species and tissues, such as oxidative damage, pro-inflammatory factors accumulation, and cellular senescence^2^. Here we investigated the impact of young ADSCs-sEVs on these markers in heart tissue. Oxidative damage is commonly viewed as a contributor to the aging process, and the heart is particularly sensitive to oxidative damage^37^. We measured MDA as a marker of lipid peroxidation^38^ and protein carbonylation as a marker of protein oxidation^39^, heart tissue of old mice exhibited higher levels of lipid peroxidation and protein carbonylation when compared to their young counterparts, indicative of increased oxidative damage in the aged heart (Fig4A-B). Remarkably, treatment with sEVs resulted in a reduction of these markers, suggesting a mitigating effect on oxidative stress-induced damage (Fig4A-B). Regarding the pro-inflammatory landscape that usually accompanies aging^40^, we measured the levels of two factors that are usually increased in aged tissues, interleukin-6 (IL-6) and interleukin-8 (IL-8), pro-inflammatory cytokines that are also tightly associated with the senescence-associated secretory phenotype (SASP)^41^. Both factors displayed an age-related increase in the heart, which was notably attenuated following sEVs treatment (Fig4C-D). As another tissue inflammatory, we measured the infiltration of T cells (CD3+) outside of the blood vessels in the heart tissue using immunofluorescence, which was almost non-existent in the young tissues and followed a similar increasing age-related pattern. Again, treatment with sEVs partially reversed this trend (Fig4E-F).

**Figure 4:**
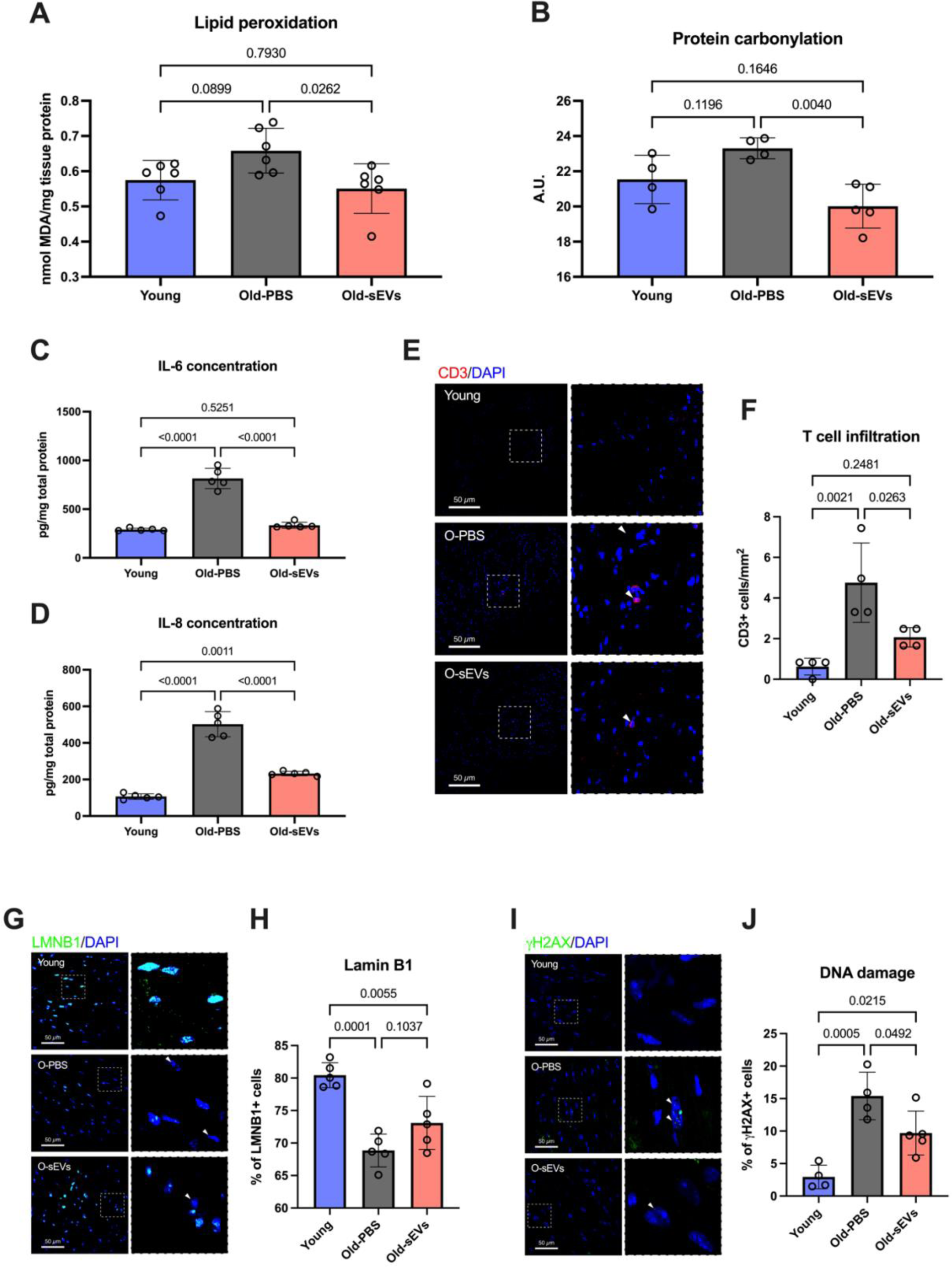
ADSC-sEVs alleviate cellular and molecular traits associated with aging in the heart of old mice. (A) Quantification of lipid peroxidation through determination of MDA levels in heart tissue. (B) Quantification of protein carbonylation as a marker of total protein oxidation in the heart tissue. (C-D) Quantification of IL-6 and IL-8 levels in heart tissue. (E-F) Representative immunofluorescence images and quantification of CD3+ cells in the heart. (G-H) Representative immunofluorescence images and quantification of LMNB1+ cells in heart tissue. (I-J) Representative immunofluorescence images and quantification of γH2AX+ cells in heart tissue. All data are shown as means ± SD. A: n=6 per group. B: n=4 for young and Old-PBS, 5 for Old-sEVs. C-D: n=5 per group. F: n=4 per group. H: n=5 per group. J: n=4 for young and Old-PBS, 5 for Old-sEVs.

We then measured two closely related markers of cellular senescence and DNA damage. We employed Lamin B1 (LMNB1) immunofluorescence to quantify cellular senescence, the loss of LMNB1 is widely used in tissues to measure the levels of senescent cells^42^. We found a reduced proportion of LMNB1+ cells in the heart with aging, while the treatment with ADSCs-sEVs showed a non-significant increase in this marker (Fig4G-H). The γH2AX marker of DNA damage exhibited an age-related increase in the heart, along with the results regarding oxidative damage to proteins and lipids, the treatment with ADSCs-sEVs reduced the levels of this marker (Fig4I-J).

These findings suggest that treatment with young ADSCs-sEVs ameliorates cellular and molecular markers associated with aging in the heart, reducing oxidative and DNA damage, and counteracting the age-related pro-inflammatory environment.

### Treatment with ADSC-sEVs switches the heart metabolome of old mice to a youthful state

Although there is a great body of evidence showing that impaired metabolism in the heart accompanies the aging process and contributes to different age-associated alterations ^43–45^, little is known about specific changes in different metabolites with age and with targeted aging interventions. To further substantiate our findings on the beneficial effects of ADSC-sEVs on aged heart tissue, we conducted a comprehensive metabolomic analysis of the heart. This allowed us to explore the alterations in the metabolic landscape of the heart that accompany aging and investigate how ADSC-sEV treatment influences these changes. Firstly, we utilized unsupervised UMAP representation on the whole metabolite set and observed a great similarity between young and old ADSC-sEVs treated mice compared to the old control mice (Fig5A, Supplementary data 1).

**Figure 5:**
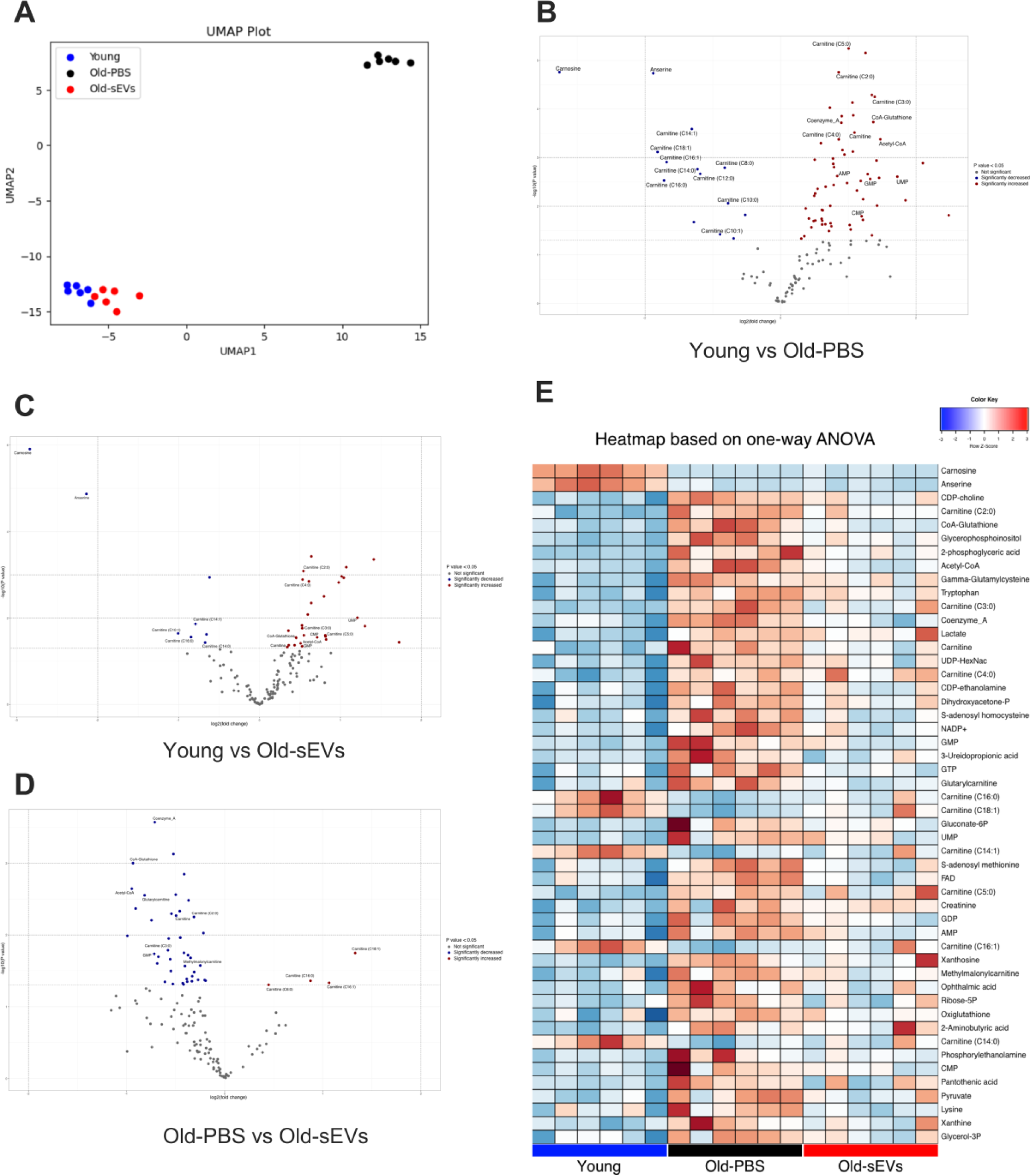
Metabolomic profiling reveals a youthful signature in ADSC-sEVs treated mice. (A) UMAP of the whole metabolite set values from heart tissue of young, Old-PBS and Old-sEVs mice. (B-D) Volcano plot showing the different metabolite levels between different comparisons. (E) Heatmap of the levels of statistically significant metabolites from ANOVA of the three groups. (F) Enrichment ratio (size of the circle) and P values (color of the circle) for the 25 most enriched metabolite sets from a metabolite set enrichment analysis (MSEA) of the significant metabolites. N=6 per group.

When comparing young and old hearts, we discovered that old mice tend to accumulate a high number of metabolites implicated in several metabolic pathways (62 upregulated and 14 downregulated in Old-PBS mice), such as Acetyl-CoA and related metabolites, GMP, AMP, CMP, or UMP, and short-chain acylcarnitines. On the contrary, young hearts showed an increased concentration of anserine, carnosine, and long-chain acylcarnitines (Fig5B). This pattern may indicate a dysregulation of mitochondrial metabolism and fatty acid oxidation, a common finding during aging^46^. Anserine is a natural derivative of carnosine, and both have been implied in cardiac health and aging, as they are important scavengers of lipid peroxidation products^47–49^. The increased acetyl-CoA levels in old mice could indicate a lower utilization or an increased production, notably, lower cytosolic levels of acetyl-CoA have been linked to increased autophagy and the beneficial effects of caloric restriction during aging ^50,51^. When comparing the heart metabolome of young and old mice treated with ADSC-sEVs we observed a similar but attenuated pattern, with a tendency to accumulate several metabolites in the old treated mice but less pronounced (30 upregulated and 8 downregulated in Old-sEVs mice), once again, young mice had higher levels of carnosine, anserine and long-chain acylcarnitines (Fig5C). Interestingly, when comparing Old-PBS and Old-sEVs mice, we found an analogous pattern, where Old-sEVs treated mice showed an increased concentration of long-chain acylcarnitines and a downregulation of several metabolites such as acetyl-CoA related (41 downregulated and 4 upregulated in Old-sEVs mice) (Fig5D).

To better visualize the changes associated with the treatment in old hearts we performed a multiple comparison analysis and represented a heatmap of the statistically significant metabolites between the three groups (Fig5E). Although it should be the task of future research, the fact that old hearts show an increased concentration of short-chain and a reduced concentration of long-chain acylcarnitines may indicate that aging influences the heart’s preference or ability to utilize different energy substrates. An increase in short-chain acylcarnitines, coupled with a decrease in long-chain acylcarnitines, might suggest a shift towards the use of medium or short-chain fatty acids for energy production.

## Discussion

The aging process is intricately linked to the decline in tissue function, and the cardiovascular system is one of the most affected. Cardiovascular diseases are the main contributors to late-life mortality in the developed world, therefore, addressing the degenerative changes associated with aging in this particular system is of great importance. Age-associated alterations of the heart tissues, including fibrosis, valvular degeneration, or cardiomyocyte hypertrophy are thought to be tightly related to molecular and cellular aspects of aging, such as inflammation, oxidative damage, or cellular senescence^52,53^.

The field of EVs research has grown exponentially in the last two decades, due to the particular ability to carry different molecules and factors between cells, making them crucial players in intercellular communication^54,55^. In the field of aging, EVs have shown potential as anti-aging factors^56,57^. We and other groups have shown some beneficial effects of EVs from different cell types in specific tissues^23,58^, such as the kidney, muscle, lungs, or the heart itself^59^.

Here we present a comprehensive study of the effect of ADSCs-sEVs on the functional, structural, molecular, and cellular parameters altered with aging in the heart of physiologically aged mice.

For the functional characterization of the heart, echocardiography assessment in mice is a complex procedure, but some established changes occur during aging in these animals, which are also common to humans, mainly an increased LV mass and reduced diastolic function, while systolic function is usually not altered^28^. ADSCs-sEVs treatment reversed the age-associated increase in LV wall thickness and the altered diastolic function reflected in the E/A wave ratio, in line with previous studies^59^. Additionally, the treadmill physical endurance test revealed beneficial sex-dependent effects of sEVs treatment, emphasizing the need for considering gender-specific responses in anti-aging interventions.

Histological analyses unveiled the impact of ADSC-sEVs on structural changes in the aged heart. Reductions in heart weight related to body weight, fibrotic tissue content, and the enhancement of vascularization collectively suggest a potential protective role of ADSC-sEVs against age-related cardiac remodeling, fibrosis, and impaired vascularization. These results support a growing body of evidence showing the potential of EVs in the modulation of fibrosis and angiogenesis^35,60^. We also provide insights into the cellular and molecular markers associated with aging in the heart. Notably, treatment with ADSC-sEVs was able to reverse the pattern of increased oxidative and DNA damage, inflammation, and cellular senescence present with aging in the mouse heart. All these processes are strongly linked, as different types of damage to the cellular integrity predispose to the acquisition of the senescent phenotype. Interestingly, cellular senescence has traditionally been viewed as a property of proliferating cells, however, the heart, as a tissue whose primary cells are post-mitotic, also shows characteristics of cellular senescence, a trait that has been termed amitosenescence^61^. The senescent phenotype also contributes to changes in the extracellular environment that lead to the accumulation of pro-inflammatory factors during aging^62^. This SASP, in turn, leads to increased cellular damage and paracrine senescence, this feedback loop is thought to contribute in a substantial way to tissue dysfunction during aging and is a target of many anti-aging interventions^63–66^. EVs, in part due to their anti-inflammatory properties^67^, are believed to counteract this loop in a senomorphic manner, shifting the extracellular environment to a non-senescent phenotype and thus reducing paracrine senescence^23,58^. Finally, we showed specific changes in a subset of metabolites that change in the aged heart. These changes include a tendency to accumulate acyl-CoA related metabolites, maybe because an inability to use different subtrates, interestingly, we showed that old hearts tend are prone to accumulate short-chain acylcarnitines and to have lower levels of long-chain acylcarnitines. In previous research, acylcarnitines levels in plasma have been associated with cardiovascular events in humans ^68^, together with our results, this may indicate a change in fatty acid metabolism in the aged heart. We also have shown how sEVs can induce a switch in the heart metabolome of old mice, resembling that of young mice. Further studies will be needed to assess whether the observed effect on the metabolism is a cause or a consequence of the beneficial effects observed in other cellular and molecular markers of aging.

Our findings add evidence to the therapeutic potential of EVs in the field of aging and cardiovascular disease as a strategy for mitigating age-related cardiac tissue alterations. However, future investigations should explore the underlying mechanisms, long-term effects, and potential translational applications of ADSC-sEVs in combating age-related cardiovascular disease.

## Acknowledgments

We would like to thank Riekelt H. Houtkooper, Bauke Schomakers, and Angelique Skantlebery for assistance in performing and/or interpreting the metabolomics.

## Funding

This work was supported by the following grants: Grants PID2020-113839RB-I00 funded by MCIN/AEI/ 10.13039/501100011033, CIAICO/2022/190 funded by Conselleria de Educació, Universitats y Ocupació and PI-2023-004 funded by PI-2023-004 to CB. Part of the equipment employed in this work has been funded by Generalitat Valenciana and co-financed with ERDF funds (OP ERDF of Comunitat Valenciana 2014-2020).

## Disclosure of interest

None of the authors declare any conflict of interest.

## Data availability statement

The datasets generated and/or analyzed during the current study are available either in the main text and supplementary files, or from the corresponding author on reasonable request. Additional data related to this paper may be requested from the authors.

## Notes

### Competing Interest Statement

The authors have declared no competing interest.

